# The spatial and temporal spread of highly pathogenic avian influenza in North America: Newton’s Cradle hypothesis

**DOI:** 10.64898/2025.12.19.695483

**Authors:** Christopher Griffin, Chiara Vanalli, Peter Hudson, Kurt Vandegrift

## Abstract

The recent emergence of highly pathogenic H5N1—especially clade 2.3.4.4b has led to widespread mortality in poultry and wild birds and has raised significant concerns for the dairy industry and human health. Migratory waterfowl are considered the main source of infection, and we used publicly available surveillance data and bird observation data from continental North America to show clear seasonal signals correlated with waterfowl movement, both on the continental scale and in three of the four flyways. In early 2024, the virus expanded its host range, and we observed a phase transition with the loss of the seasonal signal coupled with a concomitant increase in the proportion of mammalian cases. We also identified a second harmonic, with a regional east-to-west movement with infections spreading between regional flyways, followed by local viral amplification. We likened this to the movement of balls in a Newton’s Cradle with an analogy between potential and viral energy. We used bird data to identify bird species associated with viral cases and identified specific waterfowl species and highlighted the importance of predatory and scavenging birds, specifically raptors and gulls, in local amplification. These findings will help to focus surveillance strategies both at local and regional levels.

## Introduction

In recent years, avian influenza viruses (AIV) have caused significant mortality in both birds and mammals, devastated the poultry industry, affected the dairy industry, and may soon pose a major risk to human health [1–4]. Free-living waterfowl species are recognized as the natural reservoirs of low-pathogenic avian influenza viruses (LPAI), and in temperate regions, seasonal outbreaks coincide with migrating waterfowl, fueled by a large population of susceptible juvenile ducks [5–7]. Highly Pathogenic Avian Influenza (HPAI) can evolve in poultry after spillover from waterfowl, when genetic changes in the hemagglutinin (HA) gene acquire the polybasic cleavage site that allows the HA to be cleaved by a wide range of proteases, facilitating systemic infection and driving virulence [8, 9].

HPAI viruses, in particular clade 2.3.4.4b H5N1, have recently shown a large expansion in geographical distribution, extending from China, through Europe, into North and South America, and down to Antarctica [10–12]. The seasonality of human influenza has been studied in temperate regions [13–15] and analyses of links between HPAI and migrating waterfowl have also been established [16–18] such that the overarching hypothesis is that certain species of waterfowl drive the continental patterns of spillover, with local breakdowns in biosecurity leading to clusters of farm infections. However, only recently have sufficient data become available to examine this hypothesis across the four distinct flyways of North America and to examine the patterns at multiple scales. First, the continental scale, second the regional scale, where we examine patterns both within and between flyways and thirdly the local farm community scale. Here, we examine whether the HPAI cases confirmed by the National Veterinary Services Lab (NVSL) in the US and the Canadian Food Inspection Agency in Canada [19] correlate spatiotemporally with the continental migration of waterfowl species, the regional movement within and between flyways and the local amplification of the virus.

Concurrent with the geographical expansion of HPAI, there has also been an expansion in the range of hosts susceptible to 2.3.4.4b, with known infections in more than 48 mammalian species, including carnivores, pinnipeds, and dairy cattle [20–23]. Mammalian transmission of this clade was first observed in mink and foxes in Europe [24, 25]. The strain then spread to South America and moved like a wave down the west coast and then back up the east coast, causing large scale mortality of seabirds and sea mammals in 2023 [26–29]. The first recorded case of H5N1 in cattle was in February 2024 in Texas, after which it was recorded in more than 1000 farms in 17 additional states [23, 30]. Adjacent poultry farms became infected, as were cats and several species of synanthropic rodents (*Mus musculus, Rattus norvegicus*, and *Peromyscus spp*.) and more than 70 farmworkers [23, 31–34]. The maintenance of this HPAI clade in both bird and mammal populations is facilitated by the virus’s ability to bind with both avian and mammalian sialic acid receptors [20, 24, 32]. These changes in host composition of the HPAI coupled with more hosts shedding virus [35], lead us to expect an anomaly in the seasonal pattern of infection data in the spring of 2024 when the virus dramatically increased its host range.

We set out to determine whether there were detectable patterns in the spatiotemporal variation of HPAI cases and interrogated the hypothesis that migratory waterfowl drive the continental, seasonal patterns of spillover. In particular, we sought to answer the following questions: 1) Is there a coherent continental seasonal, spatiotemporal dynamic in the HPAI case data that is related to waterfowl migration? 2) Does the expansion into mammalian hosts in the spring of 2024 result in an anomaly in these seasonal data? 3) Are these patterns consistent within and between the four major North American avian flyways and do we see evidence of infection moving between flyways? 4) Can we identify key host species that drive the within and between flyway patterns that are correlated with HPAI cases?

We postulate that the signal we acquire from the continental and regional level flyways are determined by waterfowl movement but shaped by local interactions with other bird species specifically the predatory and scavenging bird species. We observe a surprisingly simple yet statistically robust double oscillator that drives HPAI dynamics within North America. In essence, the data reflect a convection-diffusion process with the bulk movement of waterfowl driving infections along the flyways, with some species showing regional movement between flyways followed by the local amplification of the virus by other, often resident species. Thus, just as the swinging balls in a Newton’s cradle transfer potential mechanical energy through a series of stationary balls, so too do the migratory waterfowl transmit viral infections to the local bird communities that amplify the virus, a double oscillator with both north-south and east-west oscillations, as illustrated in Fig. 5.

## Results

Between November 2021 and July 2025, 19,209 confirmed HPAI cases were identified. The majority were reported in the United States (86.1% wild birds, 9.8% poultry and backyard birds, and 4% mammals). Fewer than 16% were recorded in Canadian provinces (94% birds and 6% mammals). The caseload is shown in Fig. 1(a).

**Figure 1.**
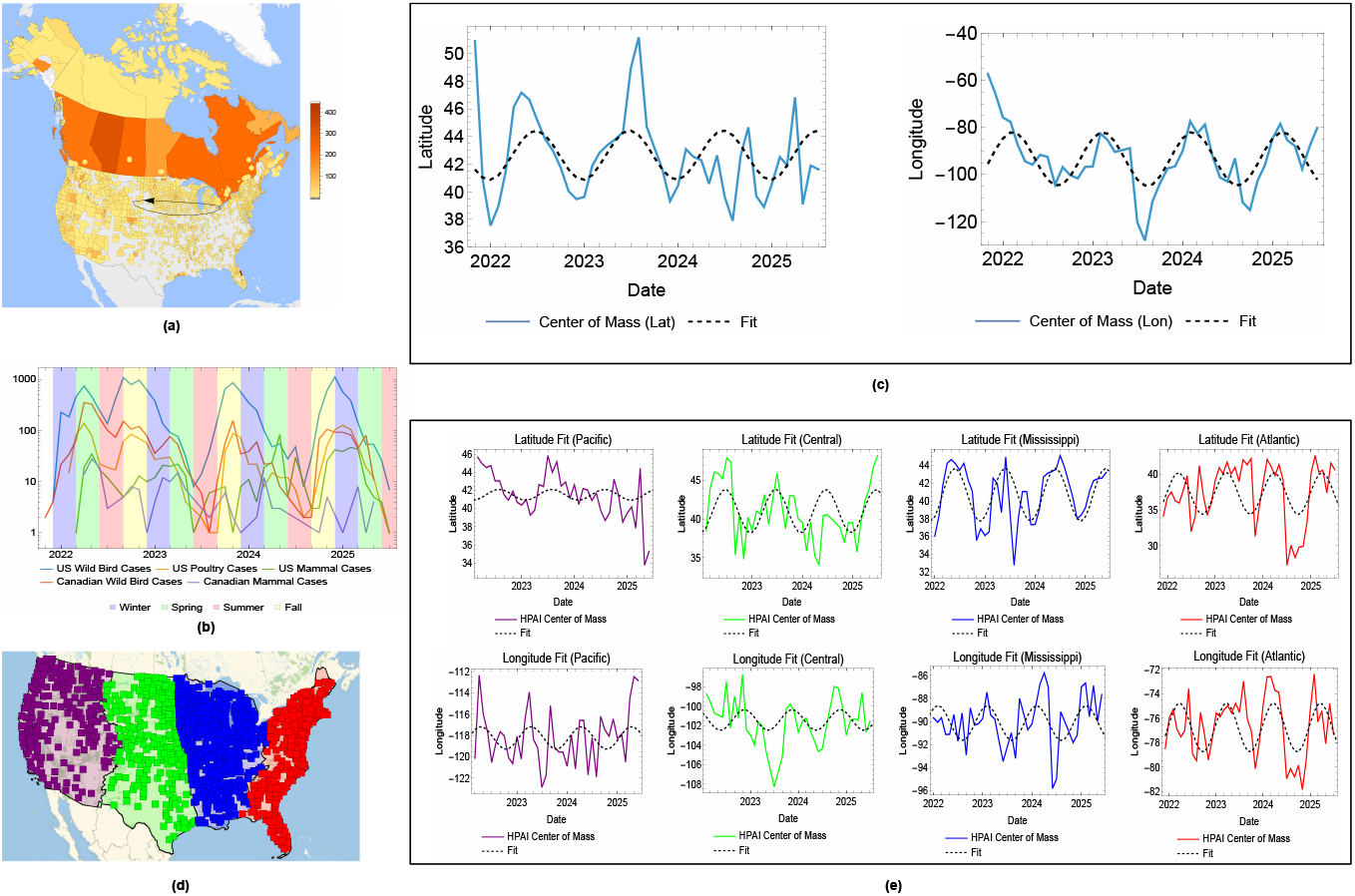
(a) Case map for the North American continent from November 2021 to July 2025 with superimposed HPAI center-of-mass ellipse. (b) Log plot of case counts over time for various HPAI data sets. (c) HPAI center of mass latitude and longitude plots with fit at the continental level. (d) Illustration of US flyways with cases mapped to flyways. (e) HPAI center of mass latitude and longitude plots with fit for each flyway.

### Spatio-Temporal Dynamics of HPAI Cases in North America

The data exhibit a clear seasonality of infection, with outbreaks coinciding with the north-south migration of birds. See Fig. 1(b). We also see oscillations in the center of mass of the HPAI cases (computation described in Methods), with annual periodicity in both the latitude and longitude of the case center of mass. See Fig. 1(c). Fits of the center of mass of infections using Eq. (1) show that the latitude of the center of mass in the continental United States (CONUS) does indeed follow the dynamics of wild bird migration, with the latitude and longitude in CONUS given by the statistically significant fits,

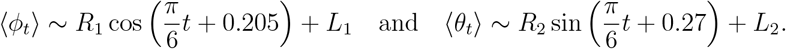

with parameter tables,

|  | Estimate | Standard Error | t-Statistic | P-Value |
| --- | --- | --- | --- | --- |
| $L_1$ | 40.07 | 0.32 | 125.3 | $1.1 \times 10^{-55}$ |
| $R_1$ | -1.52 | 0.45 | -3.40 | 0.002 |
|  | Estimate | Standard Error | t-Statistic | P-Value |
| $L_2$ | -93.96 | 1.12 | -82.47 | $4.34 \times 10^{-48}$ |
| $R_2$ | 8.65 | 1.63 | 5.32 | $3.81 \times 10^{-6}$ |

The validity of these models is confirmed at the North American continent level with modified statistically significant fits,

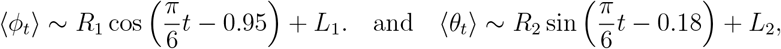

with parameter tables,

|  | Estimate | Standard Error | t-Statistic | P-Value |
| --- | --- | --- | --- | --- |
| $L_1$ | 42.65 | 0.432 | 98.73 | 0 |
| $R_1$ | -1.78 | 0.602 | -2.96 | 0.005 |
|  | Estimate | Standard Error | t-Statistic | P-Value |
| $L_2$ | -93.45 | 1.61 | -58.80 | 0 |
| $R_2$ | 11.4561 | 2.32 | 4.95 | 0.00001 |

Fits are shown in Fig. 1(c). Note that fit differences are a result of both geography and the time span of the data. See Materials and Methods for details.

### HPAI Cases across Flyways

The findings at the continental level were largely replicable within the four migratory flyways using only CONUS data. (See Fig. S3.) The center of mass of infection oscillates both latitudinally and longitudinally in all but the Pacific flyway, where no latitudinal cycle could be detected and was only weakly cyclic longitudinally (*p* = 0.06). See Fig. 1 (d) and (e). Fit details are provided in Table 1 with correlations taken between individual flyway models and the raw center of HPAI infection mass data within that flyway, as discussed in the methods section.

**Table 1.** Center of mass latitude and longitude fit properties show that there is no identifiable annual north/south pattern in data in the Pacific flyway, but all other flyways have both north-south and east-west oscillations.

| **Region** | **Correlation** | **Adjusted-$r^2$** | **Model $p$-Value** |
| --- | --- | --- | --- |
| **Latitude Models** |  |  |  |
| Pacific | 0.16 | 0 | 0.32 |
| Central | 0.51 | 0.24 | 0.0008 |
| Mississippi | 0.69 | 0.47 | $7 \times 10^{-7}$ |
| Atlantic | 0.48 | 0.22 | 0.0009 |
| **Longitude Models** |  |  |  |
| Pacific | 0.30 | 0.06 | 0.06 |
| Central | 0.33 | 0.08 | 0.04 |
| Mississippi | 0.47 | 0.2 | 0.002 |
| Atlantic | 0.58 | 0.32 | 0.00004 |

### Modeling HPAI Infection on Farms

Using only farm infections [36], a similar analysis showed no statistically significant continent-scale model for infection dynamics. That is, neither the latitude nor longitude center of mass time series showed annual oscillations using center of mass computed from HPAI cases on farms alone. However, within flyways, there were statistically significant oscillations in both latitude and longitude in the farm infection data, except (as before) in the Pacific. Fit statistics are shown in Table 2. Additional plots of the fits are provided in the SI.

**Table 2.**
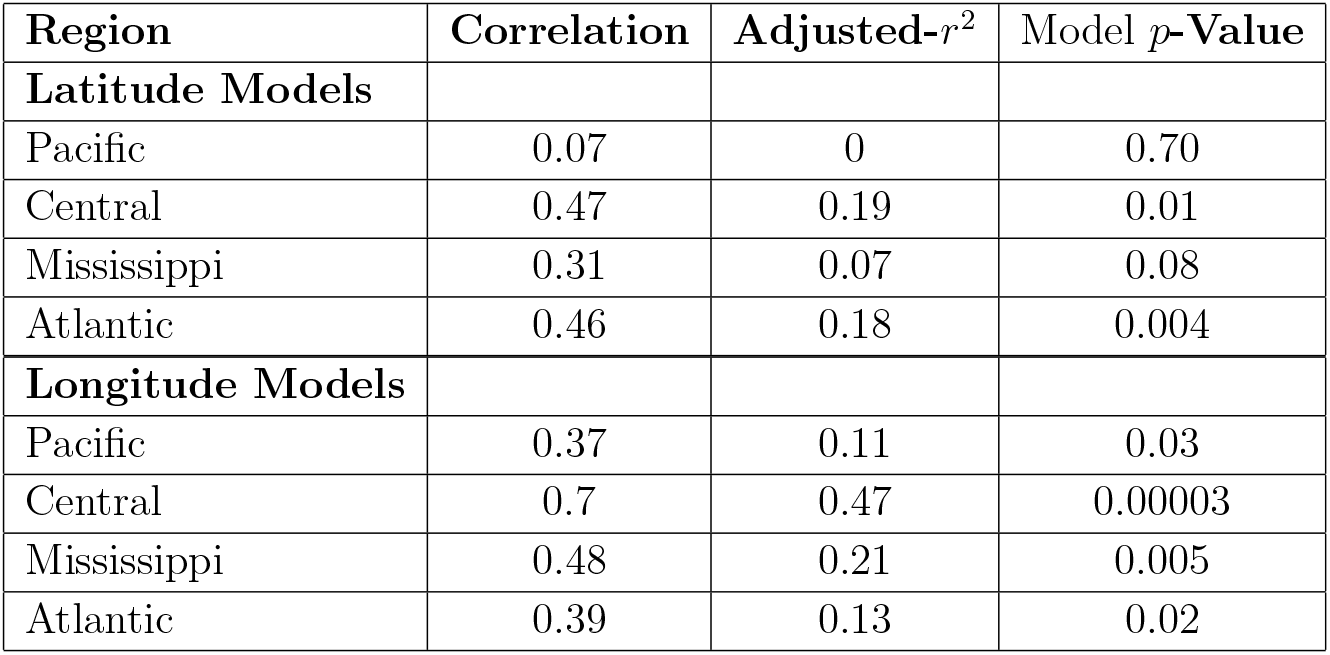
Center of mass latitude and longitude fit properties computed from farm data alone show qualitative similarity with the general flyway models using all data, with a loss of significant latitude fit in the Central flyway.

### Continent-scale Longitudinal Cycles Explained Wild Bird Movement

While we initially attempted to explain the double oscillator by weather phenomena (see SI for details), we ultimately found that the oscillation patterns could be explained by direct comparison to the centers of mass of bird species that had been identified as contracting HPAI. Of the 70 species analyzed, only 23 species showed center of mass motion that correlated with HPAI motion dynamics in both latitude and longitude (correlation above 0.3 in both latitude and longitude). See Fig. 2(a) for correlated families and Fig. 2(b) for correlated species. These 23 bird species, along with their center of mass motion and their relation to the continent-scale model, are shown in Fig. 3 (top) and this allows us to pinpoint key species that may be causing the spread of HPAI across the North American continent. Similar analysis within flyways allows us to further refine these species lists, which are shown in Fig. 3 (center). Note the role gulls play in the Pacific flyway as drivers of HPAI center of mass dynamics is similar to the role the *Anseriformes* play in the Central and Mississippi flyways. Interestingly, this pattern does not hold in the Atlantic flyway, with no species movement highly correlated with the HPAI center of mass motion in both latitude and longitude. In fact, the HPAI center of mass latitude is anticorrelated with the north-south motion of most (but not all) species. In Fig. 3 (bottom) we show species that are correlated with the HPAI model in either latitude or longitude. As noted, no species were simultaneously correlated in both.

**Figure 2.**
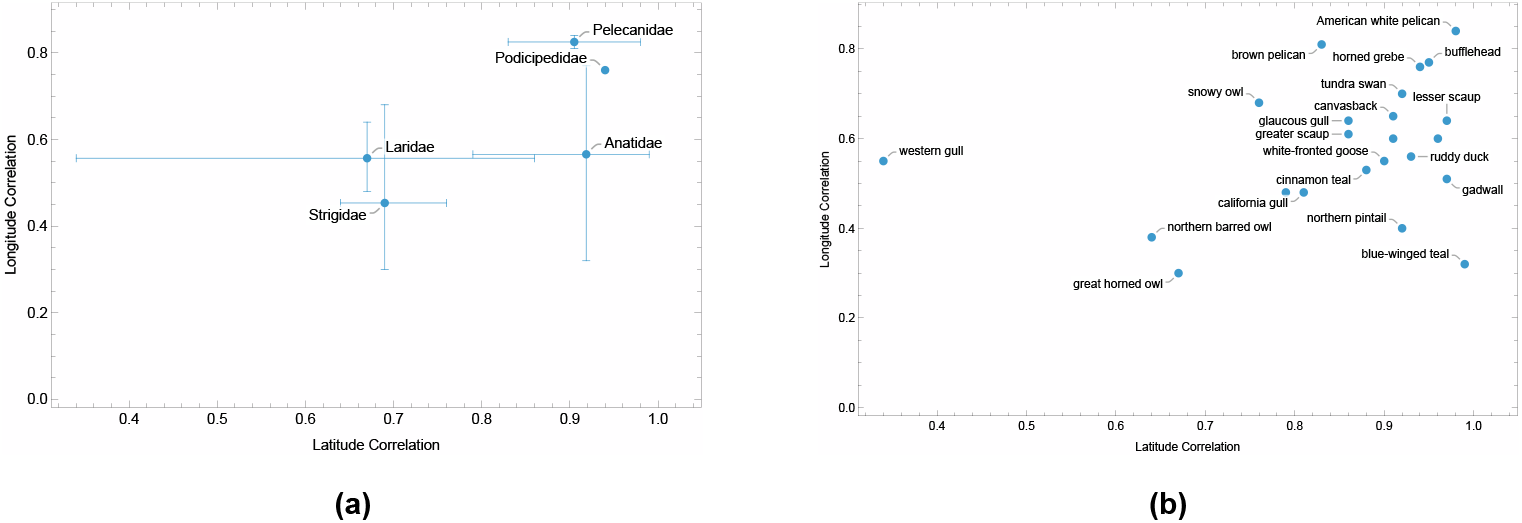
(a) Correlation-correlation plot in latitude/longitude showing bird families that are highly correlated to the HPAI motion model at the continent scale. (b) Correlation-correlation plot in latitude/longitude showing bird species that are highly correlated to the HPAI motion model at the continent scale. These data indicate which avian families and species are well positioned to influence transmission and indicates the importance of waterfowl and pelicans in driving the cases during migration at both continental and regional scales.

**Figure 3.**
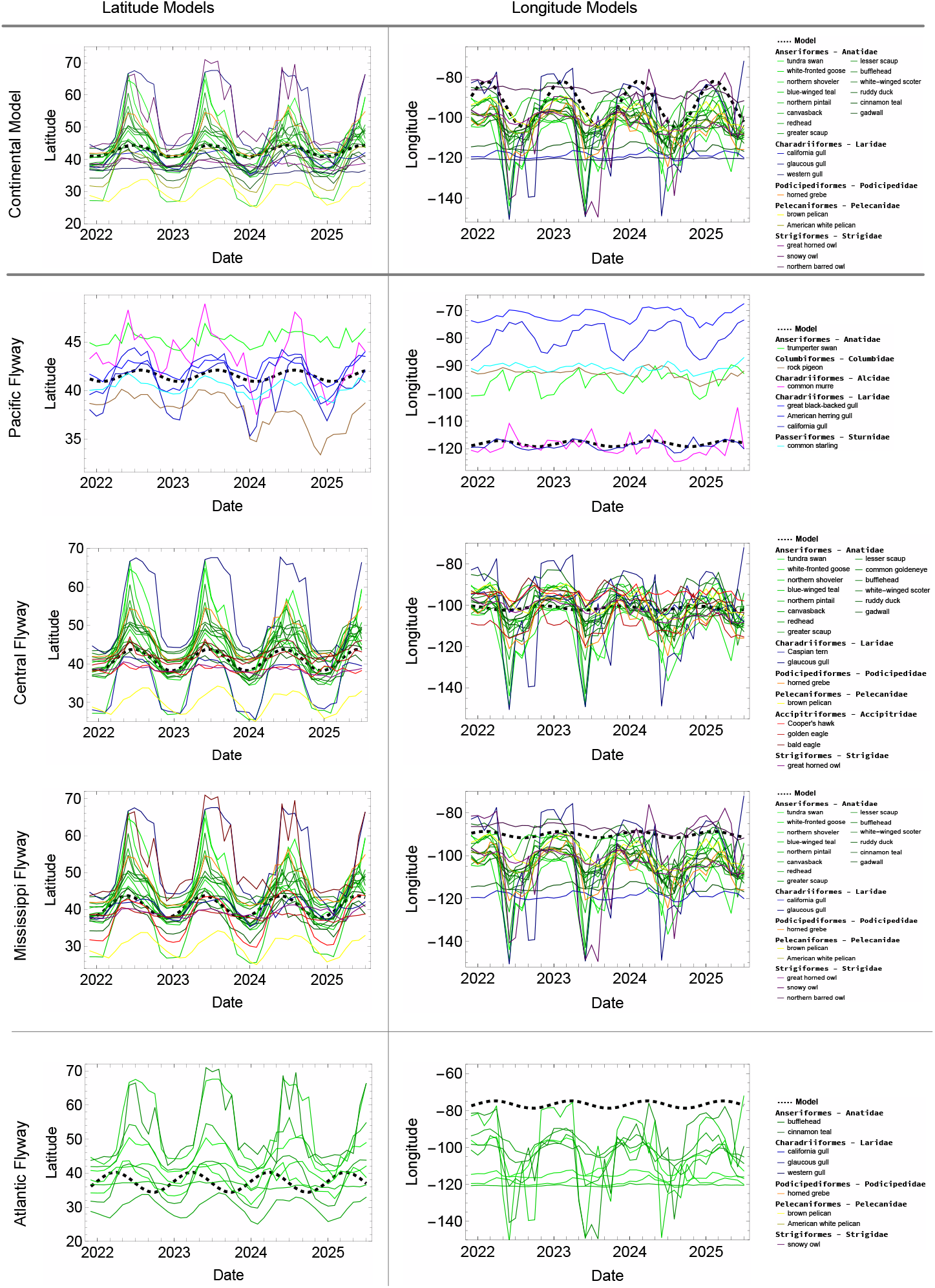
(Top) Continent-level bird species center of mass latitude/longitude positions over time and HPAI models for highly correlated bird species. (Middle) Pacific, Central, and Mississippi flyway bird species center of mass latitude/longitude over time and HPAI models for highly correlated bird species in both latitude and logitude. (Bottom) Atlantic flyway bird species center of mass latitude/longitude for species correlated to the HPAI model in either latitude or longitude. No species correlates in both in the Atlantic flyway.

### Avian Infection Networks

Analysis of monthly derived contact networks shows that the bird species identified as key hosts (see Fig. 3) are not necessarily those species that maintain HPAI infections in local areas. Degree centrality analysis suggests that local mallards, pigeons, and raptors are well positioned to play a substantial role in virus amplification and maintenance via local transmission. The number of months (out of 44) in which a species had a degree centrality in the top five in the inferred contact networks are shown in Table 3 and highlights the highly connected key species that are likely involved in sustaining HPAI infections locally. Figure Fig. 4(a) shows a single infered contact network informed by all the monthly range-based contact networks with weaker edges suppressed for clarity. (Ranges overlapped sufficiently that every species connects with several other species in at least one month.) We color and thicken the remaining edges by frequency of occurrence and adjust the size of the vertices (species) to reflect the number of times they were one of the top 5 connector species in a contact network. Colors of the vertices correspond to the number of HPAI cases observed in the data with redder vertices showing higher HPAI prevalence.

**Table 3.** Table of species ordered by number of months in which that species had a top five degree centrality.

| Common Name | Group | Highly Connected Months of 44 |
| --- | --- | --- |
| red-tailed hawk | raptor | 39 |
| mallard | waterfowl | 38 |
| ring-necked duck | waterfowl | 38 |
| peregrine falcon | raptor | 34 |
| snow goose | waterfowl | 34 |
| bald eagle | raptor | 30 |
| rock pigeon | columbid | 28 |
| great horned owl | raptor | 28 |
| greater scaup | waterfowl | 26 |
| northern pintail | waterfowl | 25 |

**Figure 4.**
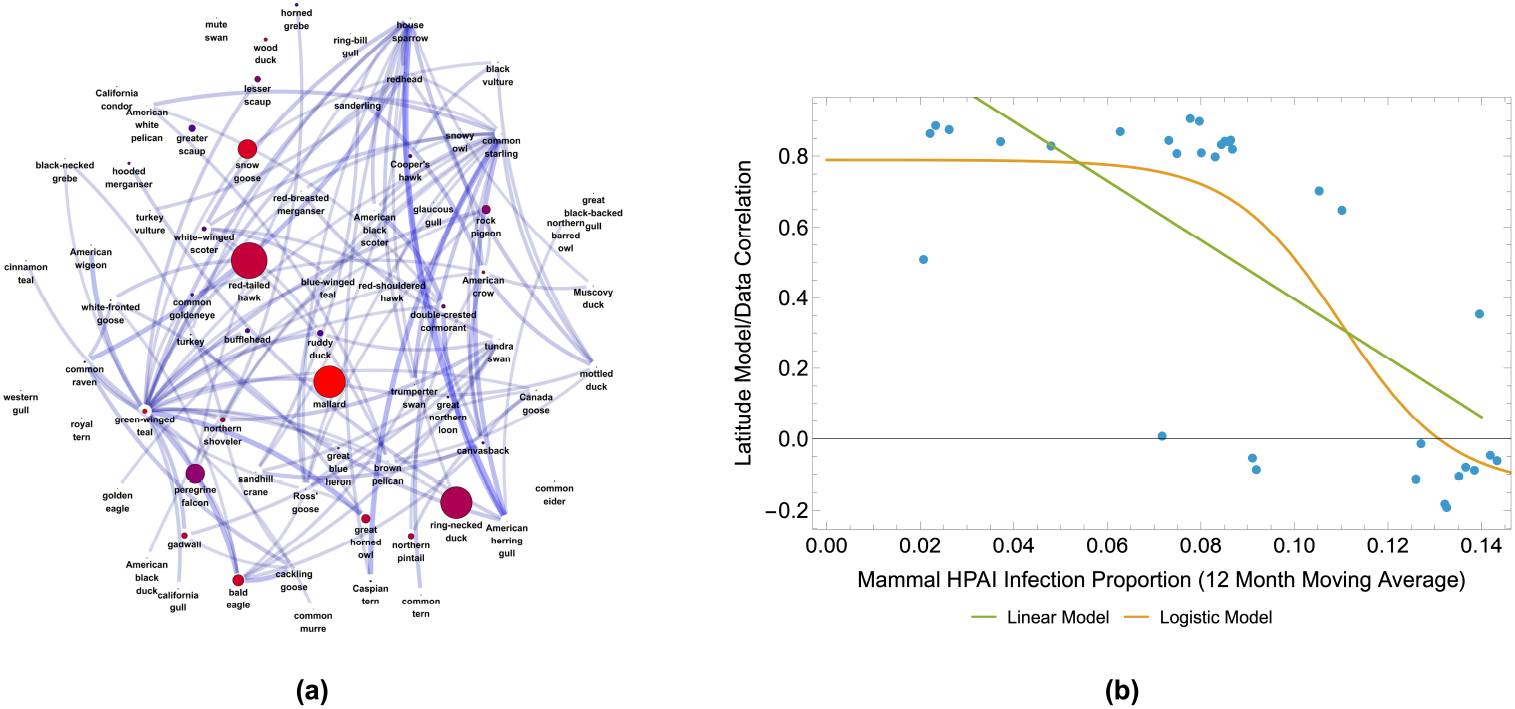
(a) A cumulative range-based contact network with edge weights proportional to the number of months two species have overlapping ranges and only edges corresponding to high overlap (> 43 months) retained. Larger nodes are more central over all months. Node coloring corresponds to the proportion of case counts in the data set. Redder vertices represent a higher number of case counts. Note that mallards and some raptors are both highly central and have a high case count, suggesting they may be connector species causing local (diffusive) spread in their environments. (b) Phase transition of correlation in latitude oscillation as a function of 12-month moving average of observed mammal HPAI infection proportion. This pattern indicates the increased host range (particularly mammals) of the virus is associated with the disintegration of the traditionally observed seasonal dynamics of HPAI.

### Latitude Model Decoherence and Mammal Infections

Examination of the HPAI latitude model and HPAI center of mass infection data suggests that the fit in the North American continent begins to decay after month 24 (approximately January 2024). See Fig. 1(c). Looking at the proportion of total HPAI cases attributable to mammals, we see a similar spike near this time. Linear and non-linear modeling show a significant relationship between the 12-month moving average of the proportion of HPAI cases attributed to mammals and the correlation (or lack thereof) between the true latitude of the HPAI center of mass (computed from the data) and the latitude model. A linear model between these two variables shows a significant negative correlation (*p* = 2 × 10^−6^) indicating that as mammals constitute a larger proportion of the infected population, the less likely one is to observe an annual oscillation in mean latitude of infection; i.e., the north-south migratory pattern is broken. See Fig. 4(b). Interestingly, the data exhibit a phase transition when approximately 10% of the HPAI cases are composed of mammals, with a logistic model given by Eq. (2), as shown in Fig. 4(b). In contrast, neither the longitudinal dynamics nor the latitude models within flyways seem to exhibit this breakdown; see Fig. 1(c) and (e).

## Discussion

### The Spatio-Temporal Dynamics of HPAI Cases in North America

Here, we examined the hypothesis that dabbling waterfowl drive the continental patterns of spillover with local breakdowns in biosecurity resulting in clusters of farm infections. Our spatiotemporal analysis of 2.3.4.4b HPAI from samples collected across North America reveals the existence of a double stochastic harmonic oscillator operating at three scales. Waterfowl first drive the continental north-south seasonal dissemination of the virus and then we identified a regional, east to west movement between flyways with probable local amplification from waterfowl by other birds. These east-west harmonics suggest a convection and diffusive epidemic dynamic that is reminiscent of Newton’s cradle. See Fig. 5. In the Newton’s cradle, the potential energy of the first ball is transferred into kinetic energy and momentum between balls, delivering energy, via the intermediate balls, to the last ball that swings up and starts the return process. In the same way, the “viral energy” in the migrating waterfowl along the eastern flyways introduces the virus to local bird populations and initiates a westward spread into neighboring flyways where it is amplified by resident birds. Our analyses indicate that this local amplification from waterfowl is mostly driven by raptors and gulls (see Fig. 5), species that are often scavengers but will also catch and consume waterfowl. These scavengers could play a very significant role in the local amplification of viruses given that HPAI can survive for 240 days in feather tissues and 160 days in muscle when kept at 4^◦^C, although viral survival decreases dramatically as temperatures increase [37]. Indeed, local temperature may play dual roles in determining both when waterbodies are free of ice and can be utilized (see SI), partially mediating migration, and providing excellent conditions for virus survival.

**Figure 5.**
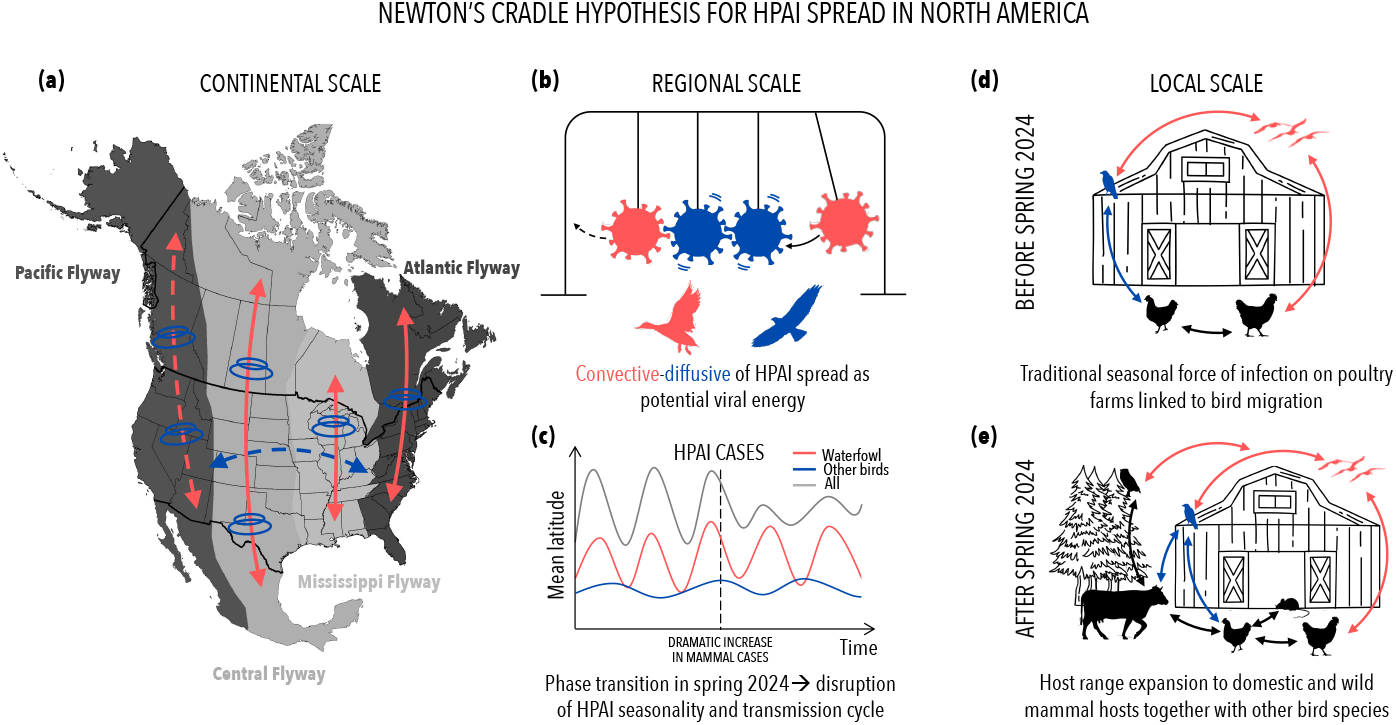
A graphical illustration of the Newton’s cradle hypothesis for the spatiotemporal spread of HPAI in North America. (a) Seasonality in the spread at the continental scale whereby waterfowl migration north and south along the flyways results in cases. This latitudinal spread for each of the four flyways is highlighted with red arrows except the Pacific flyway which is dashed and is not significant. In blue, we illustrate the regional east to west seasonal oscillation with local HPAI amplification by resident bird species. (b) The spread at the regional scale is likened to Newton’s cradle with waterfowl initiating the spread between flyways which is amplified locally by mallards and scavenging birds. (c) Graphic illustration of the temporal change in the seasonal dynamics at the continental scale after the start of 2024 when the HPAI strain extended their host range into mammals and the harmonic signature is lost. (d) Local transmission dynamics before 2024 when cases were driven primarily by waterfowl. (e) Local amplification and interspecific transmission after 2024 when HPAI extended its host range

In this analysis, we identify the species that are highly correlated both latitudinally and longitudinally with recorded cases of HPAI. It is important to note that the species selected from our analysis arise from correlations and are not based on host competence. Having said that, previous work has identified the importance of mallards as both significant amplifying hosts and long-distance dispersers of HPAI [38]. Mallards also shed high titers of HPAI virus via the oral route, without displaying overt clinical symptoms [39, 40]. Blue-winged teal and pintail have also been singled out as species involved in the maintenance and spread of HPAI [41] although more substantial experimental work is needed to elucidate the infectious period and patterns of viral shedding of these and many other species [42]. Our analysis within flyways and the inferred contact networks between bird species highlight the local importance of raptors and gulls, groups that have recently been identified as important amplifying hosts [43–45]. Of course, some of the species we identified may simply have suffered large-scale mortality and may not have played a significant role in the onward spread of infections to poultry farms or other species. For example, the Common Murre, a highly colonial pelagic seabird that has suffered large-scale mortality, is associated with HPAI outbreaks. This is not surprising given they breed in dense colonies so that infections spread rapidly through the colony producing a large local pulse of virus, which can then be transmitted to scavenging gulls and other species [46].

At a regional scale, the seasonal patterns in infection were only observed in the three flyways east of the Rocky Mountains. There was no significant pattern observed in the Pacific flyway. Although we could not demonstrate why this was the case, we suspect it related to the migration of mallards. Mallards in the Central and Mississippi flyways migrate through the country, while the Pacific mallards will often halt at rest stops, particularly when resources are available and the weather is milder with open water and little ice [47]. Interestingly, a recent phylogenetic analysis of the HPAI strains before 2022 provides evidence that about seven independent HPAI introductions to the Pacific flyway failed to spread, whereas the single introduction into the Atlantic flyway diffused westward and is the source of the ongoing outbreak [48]. Further phylogenetic evidence also shows sequences obtained in the Pacific were less likely to jump between flyways than they were from any other flyway [48].

Our initial goal was to see if we could identify the drivers of spillover from migrating birds to poultry farms, but only 9% of HPAI cases come from infected farms so the power is still insufficient to analyze this with rigor. Although we found evidence of spatio-temporal oscillations in both latitude and longitude in the flyways, suggesting correlation with migrating waterfowl, we did not see the emergent global oscillation across all flyways because of the substantially smaller sample size in farms cases. Nonetheless, the fact that HPAI oscillations in latitude and longitude are seen in the flyways, even using only farm data, suggests the same convection and diffusion action with migratory birds driving oscillations in case infection dynamics, with local birds further amplifying and diffusing HPAI and leading to continued spread. Additional data, observation, and analysis of both wild and farm birds are indicated to further test this hypothesis.

Given the emergence of HPAI 2.3.4.4b and the subsequent expansion of the viral host range into a many mammalian species, we expected to see a breakdown in the distinct seasonal pattern of cases once the infection was established. The virus was first observed in mammals in North America in 2022, and there was a marked surge in mammalian infections in March of 2024 when it infected wild carnivores, marine mammals and dairy cattle [32, 49, 50]. The data set on number of cases reflects this pattern, and we observe an increase in the proportion of infected mammals coupled with a distinct loss of latitude seasonality in the spring of 2024, see Fig. 4(b). Of course, we must be careful drawing conclusions from these data since they were not taken at random so we may be observing an increase in the sampling effort in mammals in and around infected premises. Moreover, these exhibit a proportional increase in the number of mammal cases, which could also be a consequence of a decrease in bird sampling. Nevertheless, the loss of seasonality is clear, indicating the virus had spread into the ecological community of free-living wildlife, with potential implications for seasonal farm risk and hence biosecurity planning.

### Avian Amplifiers and Networks

Using citizen science bird sightings, we identified key avian species that were associated with the movement of the center of mass in HPAI cases, both in the north-south migration at the scale of the flyways and also in the east-west movement between flyways. Interestingly, specific key species were not apparent in the Atlantic flyway. We suspect that the loss of wetland and waterfowl ponds coupled with a large population of resident mallards and Canada geese may have obscured a joint signal in seasonal oscillations because in isolation this flyway exhibits both north-south and east-west oscillations in HPAI. See Fig. 1(e).

Also interestingly, the migratory dynamics of pelicans, both the brown and the white pelican were strongly correlated with the center of mass movement of HPAI cases. They are also both indicated as important in recent phylogenetic studies [48]. Pelicans are conspicuous waterbirds, travel long distances during seasonal migrations along established flyways from breeding grounds in the northern Great Plains, Rockies, and Canada to wintering areas along the Gulf, Pacific, and Atlantic coasts. Their migration routes often overlap with those of migratory waterfowl, creating opportunities for pelicans to encounter and exchange pathogens at major stopover locations.

Our analyses also identified a number of resident species that are highly connected with the center of the infection mass and consequently are the species most likely to become exposed and could potentially play a role in the local amplification of the virus. While mallards were not identified as a species playing a large role in the north-south or east-west migration, they do become important in the local amplification. Once again, this may be a consequence of a high proportion of mallards being resident and amplifying the virus locally as opposed to moving it across the continent. We also identified the importance of several raptor and gull species. Many of these are associated with wetlands where they can either kill or scavenge on dead waterfowl and hence become exposed. These findings are again in line with the phylogenetic analysis of the virus in North America [48].

### Data Limitations

The data used in these analyses were not collected for this purpose and consequently introduce notable biases that affect the interpretation of the results. The domestic and wild animals sampled and tested for HPAI come from a wide range of sources, from free-living animals found dead, to active sampling by the USDA in and around outbreak clusters, to those sampled at wildlife rehabilitation centers and from harvested animals. Birds and mammals found dead are biased toward larger species from accessible locations. Samples collected from infected premises are more likely to be infected than samples collected at random. Likewise, harvested and wildlife at rehabilitation centers are selected samples.

Including the data from Canada greatly increased the extent of the sampling since Canada covers approximately 41% of North America, but these data were mostly at a different geographic scale than the data from the US. Most of the Canadian data were at the province level with relatively fewer at the city level. Data from the GBIF archive suffer similar geographic and taxonomic biases. These data are aggregated around highly populated areas with few data from remote areas. In North America, over 50% of birds recorded are from 100 of the most easily recognizable species, and absence is inferred simply from a lack of records. Albeit these are currently the best data available and our use of statistical mechanics reveals the continental-scale dynamics are still apparent and thus allow for a meaningful analysis.

### Conclusions

We set out to address four questions on the spatio-temporal spread of HPAI by examining the recorded cases in the US and Canada. We identified a coherent seasonal pattern in HPAI cases at the continental scale that was replicated in three of the four North American flyways (regional scale), but just as the virus expanded its host range into mammals, this pattern was disrupted, indicating that mammals became a source of infection. We also identified an interesting east-west movement of cases using this to identify movement between flyways and also the key species that are correlated with spread. Range-based contact networks inferred from GBIF data also suggest potential non-migrating amplifier hosts whose movement is not correlated with continental and regional-scale HPAI dynamics.

Our analyses support the original hypotheses that dabbling waterfowl drive the continental patterns of spillover. These analyses, coupled with a recent phylogenetic analysis of the virus, [48] lead us to make the hypothesis more refined. The parsimonious hypothesis now is that dabbling waterfowl drive the seasonal pattern of HPAI infection in all flyways except the Pacific, but these patterns are disrupted when the infection spreads into mammals. Furthermore, there is an east to west harmonic driven by waterfowl with local amplification of the virus by predatory and scavenging birds.

These findings have important consequences for surveillance. Mallards are among the most connected species and therefore remain well positioned to be key hosts for spread and maintenance within regional areas, but the significance of predatory and scavenging birds is highlighted here. We propose sampling underneath perches of raptors alongside water bodies and gull feces under roosts. There has been recent success in sampling viruses via air sampling [51], in which stratified random sampling at high-risk farms could help to identify when the virus is circulating and farms are at increased risk. Concomitant sampling alongside outbreaks can strengthen the power and robustness of phylodynamic approaches.

## Materials and Methods

### Data Overview

HPAI case data for wild birds [52], commercial and backyard flocks [36] and mammals [31] were obtained from both the Animal and Plant Health Inspection Service (APHIS) arm of the United States Department of Agriculture (USDA) and the Canadian Food Inspection Agency [19] (accessed on August 9, 2025). The time series data began in December 2021. Case counts (rather than animal counts) were used to normalize size comparisons due to extraordinarily large outbreaks, primarily in laying hen operations. For example, an outbreak at a poultry farm in Lancaster, PA is treated as a single case even though it may involve millions of co-located birds. The data was then aggregated into case counts per month. Data from Hawaii were removed because it is geographically isolated. The dates and county/city/province names were automatically interpreted as date objects and geographic entities via Mathematica. Counties, provinces, or cities were converted into latitude and longitude using Wolfram’s knowledge base, using the coordinates of the geographic center of the administrative unit. Interpretation or conversion failures were dropped or corrected by a secondary algorithm provided in the code. In total, there were 19, 209 cases in the data set. Of those cases, 18, 744 were successfully associated with geographic coordinates and specific months and are used in this analysis. To determine if the patterns observed at the continental scale data were also evident in the four flyways within the continental US, ESRI shapefiles for flyways were obtained from the United States Fish and Wildlife Service [53]. We used approximately 70*M* citizen scientist reported bird locations provided by the Global Biodiversity Information Facility [54] for the 70 species that had at least ten reported avian flu cases as given in the data from the US Department of Agriculture [31, 36, 52]. The species are listed in the SI.

### HPAI Motion Dynamics

Our analysis of HPAI dynamics was inspired by methods from statistical mechanics with the assumption that microscopic (bird) interactions produce mesoscopic (species) and macroscopic (continent or flyway) bulk dynamics. To identify viral flow dynamics in birds at the level of the continental United States (CONUS) and the North American continent, we removed all mammal data and considered only data from wild birds and the commercial and backyard birds within the USDA and Canadian data sets. When considering only CONUS, data from Canada, Alaska were removed. Data from Hawaii were removed in all cases. The data set has the form,

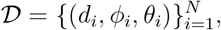

where *d*_*i*_ is the date of the *i*^th^ observation, which occurs at latitude *ϕ*_*i*_, and longitude *θ*_*i*_. Within each month, cases were grouped by latitude and longitude so that in month *t* we have a data set of the form,

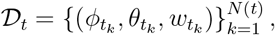

where 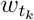 is the number of cases occurring at coordinate 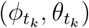 in the given month *t*, which has *N* (*t*) total cases.

In any given month, there may be HPAI cases spread throughout the United States, however we extract bulk infection dynamics by examining the time-varying center of mass of the infection given by,

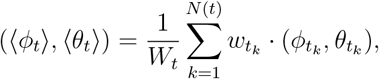

where,

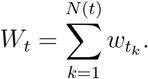

We used a similar construction when building centers of mass for the various species of wild birds included in our analysis.

We used the following functional forms as ansatz for the HPAI infection centers of mass,

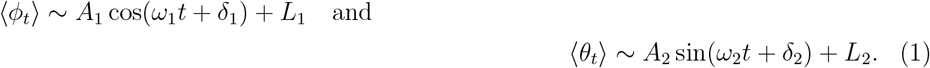

We performed initial nonlinear fitting, constraining 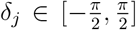. From this, we fixed *ω*_*i*_ = *π/*6, corresponding to a 12 month periodicity (as confirmed by the initial fit, discussed in the results section). We then varied *δ*_*j*_ and used a best fit multiple linear model approach, to estimate *δ*_*j*_, and to fit *L*_*j*_ and *R*_*j*_.

For analysis by flyway, we divided the data along the longitudinal span of the continental United States into the four migratory flyways [55] using ESRI shapefiles. For each flyway, we performed the same analysis as described above, with nominally modified ansatz, as described in the SI.

### Analysis of Mammal Impact

To study the impact of mammal infections on HPAI dynamics, we computed the proportion of observed HPAI cases that were attributed to mammals, denoted 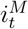. We compared the 12-month moving average of 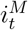 (denoted 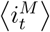) and compared it to 12-, month correlations between ⟨*ϕ*⟩, the latitudinal model, and the raw center of mass data. We denote these correlations as 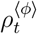. In addition to building a standard linear model relating 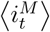 and 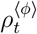, we also fit the logistic model

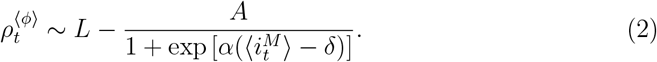

As with the nonlinear fits for latitude and longitude centers of mass, we used a parameter space search to find *α* and *δ* by minimizing the sum of squares and used a standard (linear) fit to find *A* and *L*.

### Range-Based Contact Networks

To construct speculative contact networks, we used empirical range re-construction by assuming that monthly bird observations were normally distributed within a local region. The resulting distances from the center of mass (of the each bird species in each month) is then (approximately) a *χ*^2^ random variable (see SI for details). Using this observation, we created a (spherical) circle around each species center of mass (in each month) with a radius equal to the mean distance from the center of mass to true bird location. We assume this geographic region approximates an area in which sufficient contact can be maintained to ensure viral propagation. An edge was assigned between two species in a given month if their inferred infection ranges overlapped. We used a standard degree centrality [56] on these networks. To construct Fig. 4(a) we joined all networks together, suppressing edges that were low-weight and coloring vertices as discussed in the main body of the text.

## Acknowledgement

This material is based upon work that is supported by the Animal and Plant Health Inspection Services, United States Department of Agriculture, under award number AP23OA000000C025. Kurt Vandegrift is supported by Pennsylvania’s Center for Poultry and Livestock Excellence (Award# CPLE23-11).

## Notes

https://www.aphis.usda.gov/livestock-poultry-disease/avian/avian-influenza/hpai-detections/mammals

https://www.aphis.usda.gov/livestock-poultry-disease/avian/avian-influenza/hpai-detections/commercial-backyard-flocks

https://www.aphis.usda.gov/livestock-poultry-disease/avian/avian-influenza/hpai-detections/wild-birds

https://www.fws.gov/partner/migratory-bird-program-administrative-flyways

https://cfia-ncr.maps.arcgis.com/apps/dashboards/89c779e98cdf492c899df23e1c38fdbc

https://www.aphis.usda.gov/livestock-poultry-disease/avian/avian-influenza/hpai-detections/hpai-confirmed-cases-livestock

https://www.aphis.usda.gov/livestock-poultry-disease/avian/avian-influenza/hpai-detections/mammals

### Competing Interest Statement

The authors have declared no competing interest.

