## Supplemental Info for "The spatial and temporal spread of highly pathogenic avian influenza in North America: Newton’s Cradle hypothesis"

### The spatial and temporal spread of HPAI in North America: Newton's Cradle Hypothesis for Spread Across the Continent - SI

**Christopher Griffin[1,*], Chiara Vanalli[2], Peter Hudson[3] and Kurt Vandegrift[3,*]**

[1]Applied Research Laboratory, The Pennsylvania State University, University Park, PA 16802
[2]Environmental Computational Science and Earth Observation Laboratory, École Polytechnique Fédérale de Lausanne, Sion 1950, Switzerland
The Center for Infectious Disease Dynamics, Department of Biology and Huck Institutes of the Life Sciences, The Pennsylvania State University, University Park, PA 16802
[*]Corresponding Authors

#### Additional Details on Mammal Case Distribution

In Fig. 1 we show the yearly case count and compare it to the yearly case count of mammals. There is a clear increase in mammal cases in 2023 and a clear spike in all cases in 2022. The distribution of mammal case counts is shown in Fig. 2 indicating a skew in the data for mammals toward the Pacific and Central flyways, except in Canada, where the data is more uniformly distributed longitudinally.

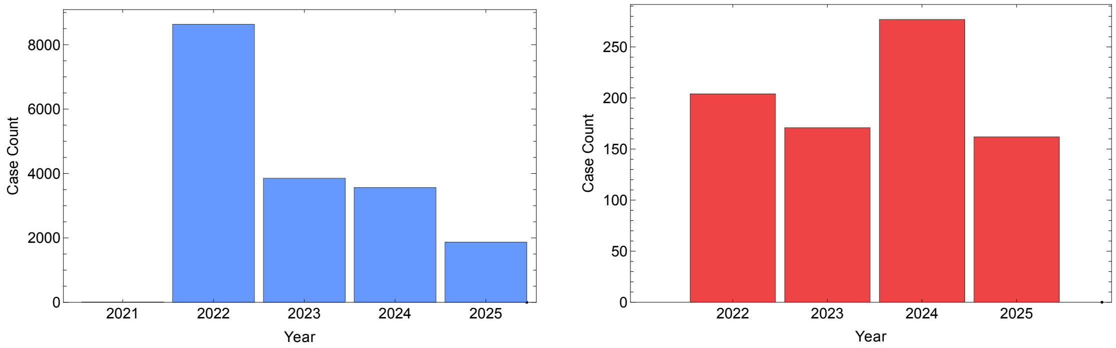


**Figure 1.** (Left) Total case count by year showing consistent decreasing case loads year-on-year. (Right) Total mammal case count by year showing an increase in 2024.

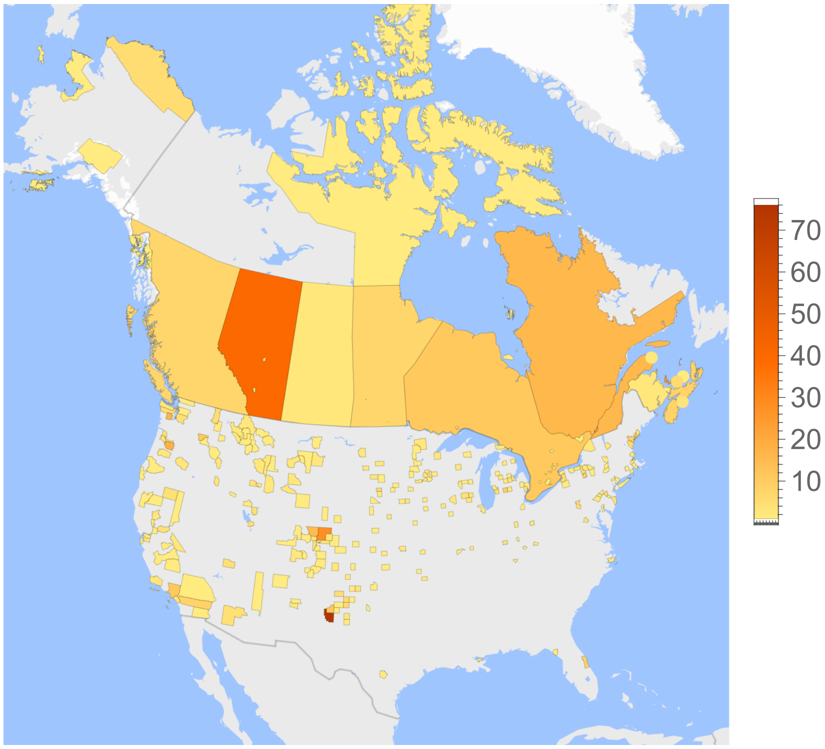


**Figure 2.** A geohistogram showing the distribution of mammal cases with data skewed toward the Pacific and Central flyways in the United States.

#### Additional Details on Flyway and Farm Fits

We divided the longitudinal span of the continental United States into the four migratory flyways [1] using ESRI shapefiles provided by the United States Fish and Wildlife Service [2]. Flyway time-series are shown in Fig. 3. Interconnectivity between the flyways, as shown by [3] suggests that HPAI may spread between these routes. Analysis shows similar oscillatory patterns within the flyways with caveats.

For each flyway, we fit the latitude and longitude ansatz,

$$\langle \phi^f \rangle \sim R_1^f \cos\left(\frac{\pi}{6}t - \delta_1^f\right) + L_1^f \quad \text{and} \quad \langle \theta^f \rangle \sim R_2^f \sin\left(\frac{\pi}{6}t - \delta_2^f\right) + L_2^f,$$

to the time-varying infection center of mass within each region. Here $f \in \{1,2,3,4\}$ correspond to the four regions numbered from west to east. The resulting centers of mass and fits are shown in Fig. 1 (e) of the main text. Quality-of-fit metrics are given in Table 1 of the main text. Fits for HPAI infections using only farm infections within flyways are shown in Fig. 4. Quality-of-fit metrics are given in Table 1 of the main text. A comparison of the HPAI latitude and longitude models in the continental United States using all bird cases and those models using just farm cases is shown in Fig. 5. We have included the Pacific flyway, despite its general lack of statistically significant models. Notice in both the Central and Mississippi flyways, the all-bird HPAI model leads the farm model, suggesting (as would be expected) that HPAI infections are being seeded by migratory birds. The opposite is true in the Pacific and Atlantic flyways. As noted, the Pacific flyway did not have a statistically significant latitude or longitude oscillation. The Atlantic flyway, is highly anomalous, with a general lack of fit for HPAI to migration patterns. This explains the fact that HPAI on farms seems to precede HPAI in general and further illustrates the Atlantic flyway anomaly discussed in the main text.

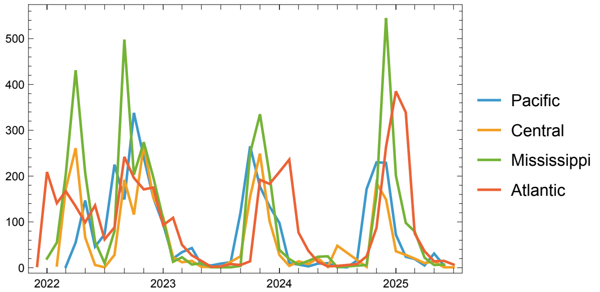


**Figure 3.** Time series of HPAI infections within the four North American flyways showing variation in seasonal infection duration and start times.

#### HPAI and Temperature: Data Processsing and Modeling

To determine whether HPAI and bird migration (longitudinally and latitudinally) follow a temperature isocline, we retrieve the average monthly temperatures from November

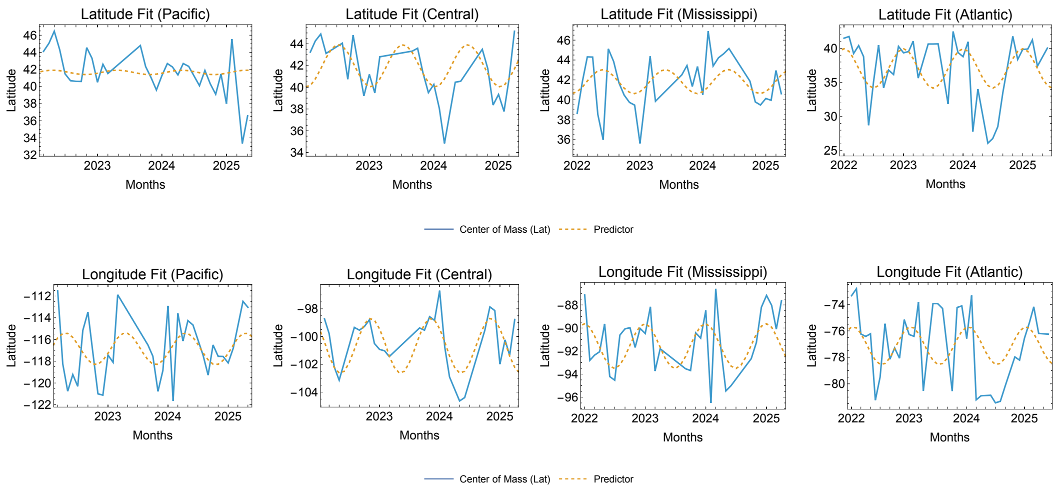


**Figure 4.** (Top) Fit of farm HPAI center of mass latitude variation within flyways. (Bottom) Fit of farm HPAI center of mass longitude variation within flyways.

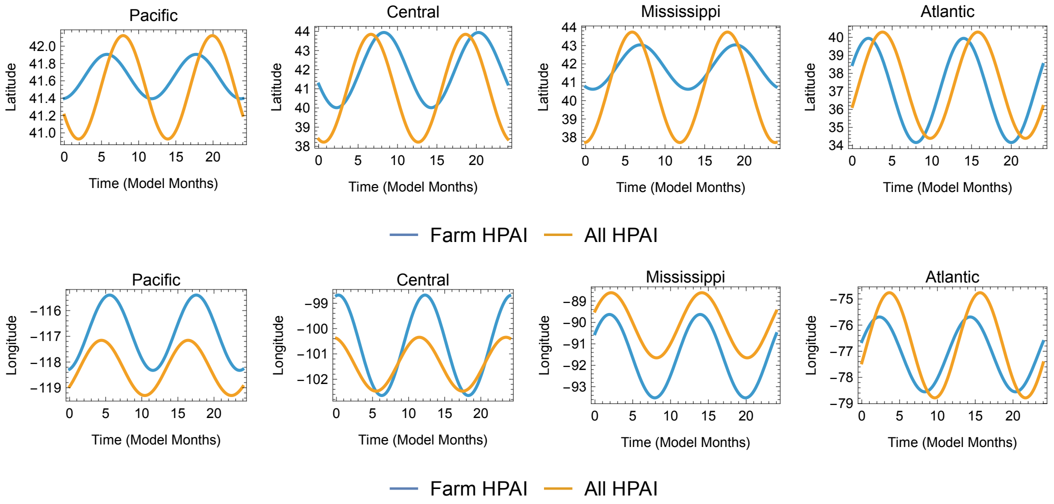


**Figure 5.** (Top) Comparison of farm HPAI latitude model and all bird HPAI latitude model in the continental United States. (Bottom) Comparison of farm HPAI longitude model and all bird HPAI longitude model in the continental United States.

2021 through July 2025, the period of investigation from Wolfram's Knowledgebase. We used a 1 degree grid of latitude and longitude to create sample points $\{(\Phi_i, \Theta_i)\}_{i=1}^{M}$, with $\phi \in [25, 80]$ (degrees) and $\theta \in [-130, -70]$ (degrees) so that $M = 3,416$. At each grid point $i$, let $T_{i\tau}$ be the average monthly temperature in month $\tau$. If $\tau = 0$ indicates November 2021, then $\tau$ is an integer from 0 to 44. Define the indicator,

$$Z_{i\tau} = \begin{cases} 1 & \text{if } T_{i\tau} > \bar{T} \\ 0 & \text{otherwise.} \end{cases}$$

This indicator determines whether a grid point has an average monthly temperature above the thermal threshold $\bar{T}$. We test two thresholds: i)$\bar{T} = 0°$C following the freezing temperature hypothesis suggested by [4]; and ii) $\bar{T} = 7°C$ to ensure that the minimum temperature would not reach 0°C, avoiding standing water freezing; for this, the mean must be higher than the local diurnal swing, which ranges from about 4°C in coastal, humid regions to 7°C in the more dry inland regions. The mean latitude and longitude above $\bar{T}$ are given by,

$$\langle \Phi_\tau \rangle = \frac{1}{N_\tau} \sum_i \Phi_{i\tau} Z_{i\tau} \quad \text{and} \quad \langle \Theta_\tau \rangle = \frac{1}{N_\tau} \sum_i \Theta_{i\tau} Z_{i\tau},$$

where,

$$N_\tau = \sum_i Z_{i\tau},$$

is the total number of grid points with monthly average temperature above 7°C in month $\tau$. The mean latitude and longitude above 7°C in month $\tau$ is shown in Fig. 6. Notice the strong oscillation in latitude (as expected) and the weak oscillation in longitude.

We now compare these latitude and longitude dynamics to the models for infection mean latitude and longitude, shown in Fig. 7. Visually, the mean infection latitude over time appears highly correlated with the mean latitude at which the monthly average temperature is above 7°C. See Fig. 7 (left). In particular, the correlation between these time series is $\rho_\Phi = 0.96$, proving conclusive support for the hypothesis that HPAI are following migration patterns, which are known to follow thermoclines, until they are disrupted in 2024 (see main text).

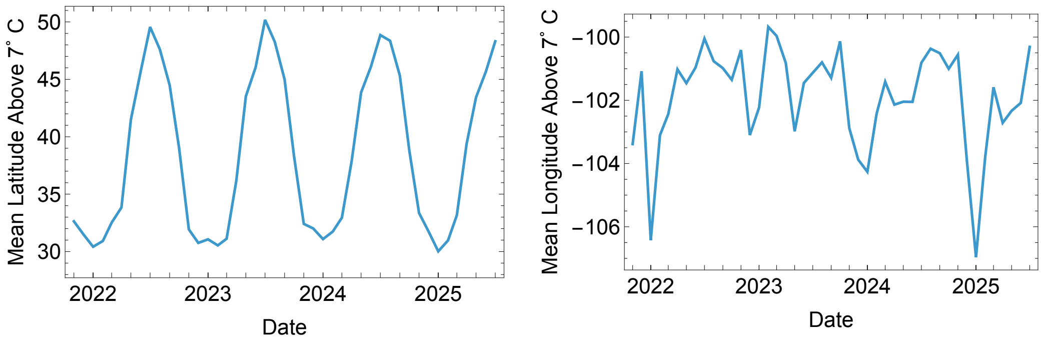


**Figure 6.** (Left) Spatial average of latitudes above 7°C with monthly average temperature above 7°C from November 2021 through July, 2025. (Right) Spatial average of latitudes with monthly average temperature above 7°C from November 2021 through July 2025.

Interestingly, the mean longitude of infection is out of phase with the mean longitude above 7°C by approximately six months; see Fig. 7 (right). The correlation between these time series is $\rho_\Theta = -0.47$, suggesting the longitudinal temperature gradient may be causing the east-west oscillation, but through an unusual mechanism, which is identified through GBIF data analysis (see main text). In exploring this potential mechanism, we found that the center of mass for the 70 bird species (see

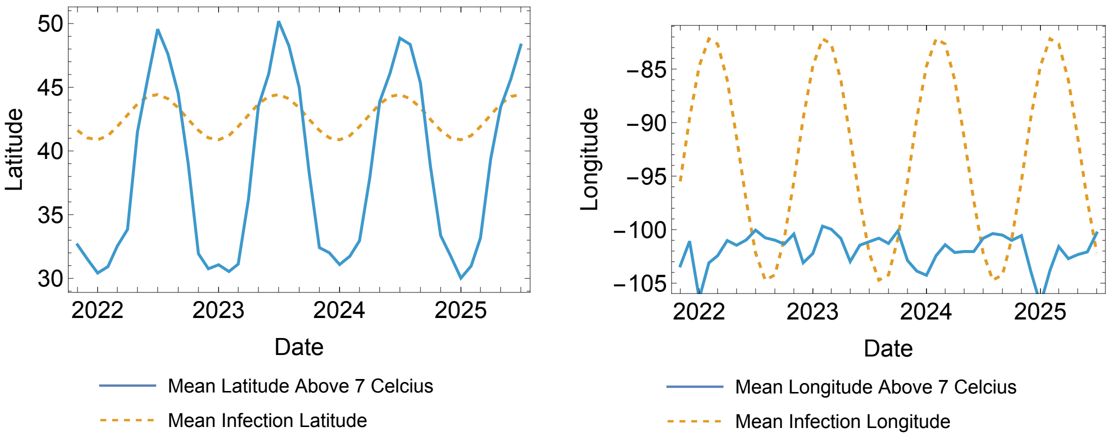


**Figure 7.** (Left) Mean latitude of infection and mean latitude at which the average monthly temperature is above 7°C show a high degree of correlation ($\rho_\Phi = 0.96$). (Right) Mean longitude of infection and mean longitude at which the average monthly temperature is above 7°C show correlation with a six-month offset ($\rho_\Theta = -0.82$).

Methods, main text) correlates well with both the mean latitude and longitude above 7°C, as expected [5–7] (latitude correlation was 0.86, longitude correlation was 0.43). However, as noted in the main text, 23 species showed center of mass motion that correlated with HPAI motion dynamics in both latitude and longitude (correlation above 0.3 in both latitude and longitude) and were therefore correlated with the mean latitude above 7°C but were anti-correlated with the mean longitude above 7°C.

#### Additional Details on GBIF Data

To more thoroughly explain the longitudinal oscillation, we used $\sim 70M$ citizen scientist-reported locations provided by the Global Biodiversity Information Facility [8] for the 70 species that had at least ten reported avian flu cases as given in the data from the US Department of Agriculture [9–11]. The species are listed in Table 1. For species $s$, we constructed a monthly center of mass $(\langle \phi_t^s \rangle, \langle \theta_t^s \rangle)$. These were used as raw data and compared to the continent-scale HPAI model. We also constructed a global bird center of mass $(\langle \phi_t^G \rangle, \langle \theta_t^G \rangle)$ and analyzed this as a control.

Using the center of mass of all species, we see that the longitude correlates well with the mean longitude above 7°C (the correlation is 0.43), suggesting that average bird motion is correlated to weather conditions both latitudinally and longitudinally. This is shown in Fig. 8 and suggests that HPAI spread is not being caused by the "average bird" but by some subset of birds whose migration patterns are distinct from the average.

Fig. 9 shows a graphic representation of HPAI dynamics over space and time with the 23 correlated species as well as the center of mass model for the North American continent. The figure shows the clear north/south and east/west motion of HPAI over time and its correlation with specific bird species. A movie showing this is also provided.

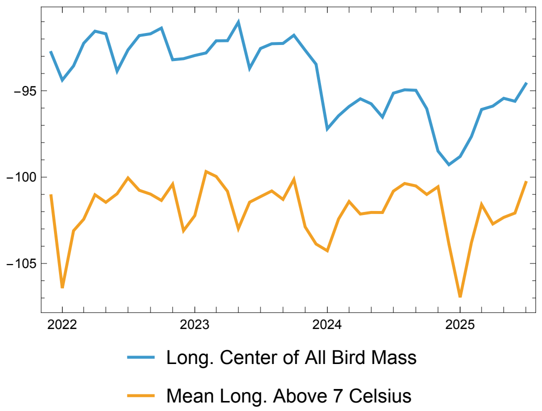


**Figure 8.** A comparison of the center of mass of all birds in the GBIF sample and the mean longitude above 7°C.

#### Additional Details on Contact Network Reconstruction

We can use empirical range reconstruction to approximate contact networks among birds and hypothesize which species may be acting as key hosts. To do this, we assumed that monthly bird observations were normally distributed within a local region. If $(\langle \phi_t^s \rangle, \langle \theta_t^s \rangle$ is the center of mass (the maximum likelihood estimator of the mean of the normal distribution), then the distances of the observations from that mean, denoted, $d_{t_i}^s$ are (approximately) scaled $\chi^2$ distributions. The approximation is not precise because the normality assumption may be only partially correct for some species, and distance must be taken on the sphere, a manifold, not a Euclidean space. Nevertheless, for a two-dimensional Euclidean space, we would expect,

$$r_t^s = \frac{\langle d_t^s \rangle}{S_t^s} \approx 2,$$

the parameter of the $\chi^2$ distribution, where $\langle d_t^s \rangle$ denotes the mean distance from the center of mass and $S_t^s$ denotes the standard deviation of the distances. A histogram of this fraction over all months and species is shown in Fig. 10. Notice the ratios cluster around the mean $\langle r \rangle = 2.3 \approx 2$, as expected. Thus, we use the mean distance to approximate a range in which a bird species has sufficient mass to create a non-trivial contact rate with other fauna. That is, the infective range for species $s$ in month $t$ is a (spherical) circle of radius $\langle d_t^s \rangle$.

For each month, we created a contact network with an edge between two species if their approximated ranges (the circles) overlapped. An example graph for October 2023 is shown in Fig. 4(a) of the main text. Using these networks, we computed the degree centrality for each species (vertex) in the network and then found those most highly connected species in each month. This was done by finding the highest five degree values and including all species with these degree values, including ties. These results are given in Table 3 of the main text.

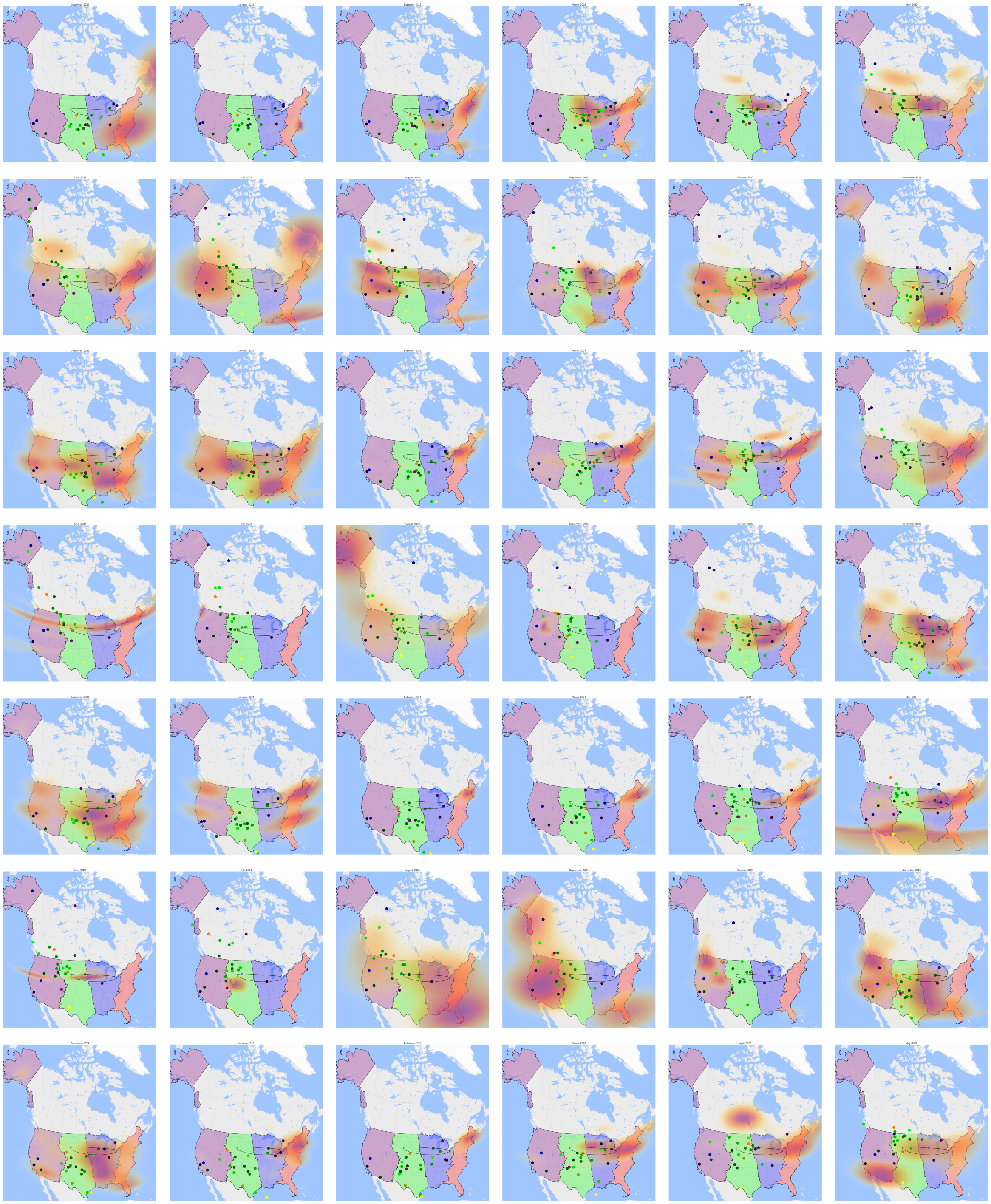

**Figure 9.** Visualization of the infection density over time, its center of mass and estimated center of mass. (Hexagons are the geo-histogram bins, colored circles are bird species, the blue square with black track shows the continental level HPAI center of mass model.

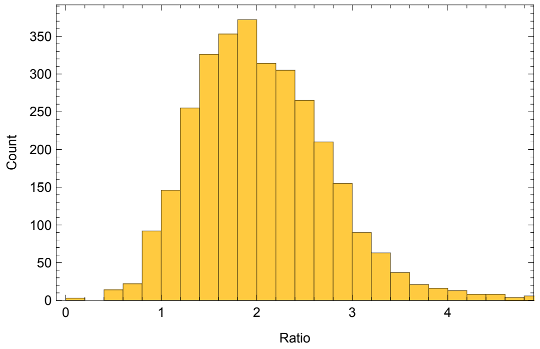


**Figure 10.** Histogram of the ratio of the mean bird distance from the center of mass over all months and species.

livestock-poultry-disease/avian/avian-influenza/hpai-detections/commercial-backyard-flocks

| **Species Name** | **Common Name** | **Species Name** | **Common Name** |
|---|---|---|---|
| *Bubo scandiacus* | snowy owl | *Thalasseus maximus* | royal tern |
| *Accipiter cooperii* | Cooper's hawk | *Aythya valisineria* | canvasback |
| *Anas carolinensis* | green-winged teal | *Mareca americana* | American wigeon |
| *Sterna hirundo* | common tern | *Bucephala clangula* | common goldeneye |
| *Cygnus columbianus* | tundra swan | *Spatula cyanoptera* | cinnamon teal |
| *Larus hyperboreus* | glaucous gull | *Anser caerulescens* | snow goose |
| *Anser rossii* | Ross's goose | *Mareca strepera* | gadwall |
| *Branta hutchinsii* | cackling goose | *Larus occidentalis* | western gull |
| *Podiceps auritus* | horned grebe | *Pelecanus erythrorhynchos* | American white pelican |
| *Anser albifrons* | white-fronted goose | *Grus canadensis* | sandhill crane |
| *Melanitta deglandi* | white-winged scoter | *Lophodytes cucullatus* | hooded merganser |
| *Anas fulvigula* | mottled duck | *Bubo virginianus* | great horned owl |
| *Mergus serrator* | red-breasted merganser | *Podiceps nigricollis* | black-necked grebe |
| *Gymnogyps californianus* | California condor | *Coragyps atratus* | black vulture |
| *Anas acuta* | northern pintail | *Larus smithsonianus* | American herring gull |
| *Aythya marila* | greater scaup | *Larus delawarensis* | ring-bill gull |
| *Aythya americana* | redhead | *Strix varia* | northern barred owl |
| *Uria aalge* | common murre | *Corvus corax* | common raven |
| *Spatula clypeata* | northern shoveler | *Oxyura jamaicensis* | ruddy duck |
| *Somateria mollissima* | common eider | *Corvus brachyrhynchos* | American crow |
| *Melanitta americana* | American black scoter | *Cathartes aura* | turkey vulture |
| *Aquila chrysaetos* | golden eagle | *Aix sponsa* | wood duck |
| *Hydroprogne caspia* | Caspian tern | *Phalacrocorax auritus* | double-crested cormorant |
| *Aythya affinis* | lesser scaup | *Sturnus vulgaris* | common starling |
| *Falco peregrinus* | peregrine falcon | *Anas platyrhynchos* | mallard |
| *Larus marinus* | great black-backed gull | *Buteo lineatus* | red-shouldered hawk |
| *Anas rubripes* | American black duck | *Cygnus buccinator* | trumpeter swan |
| *Bucephala albeola* | bufflehead | *Pelecanus occidentalis* | brown pelican |
| *Cairina moschata* | Muscovy duck | *Haliaeetus leucocephalus* | bald eagle |
| *Calidris alba* | sanderling | *Buteo jamaicensis* | red-tailed hawk |
| *Aythya collaris* | ring-necked duck | *Passer domesticus* | house sparrow |
| *Gavia immer* | great northern loon | *Columba livia* | rock pigeon |
| *Cygnus olor* | mute swan | *Meleagris gallopavo* | turkey |
| *Spatula discors* | blue-winged teal | *Branta canadensis* | Canada goose |
| *Larus californicus* | California gull | *Ardea herodias* | great blue heron |

**Table 1.** Table of species data extracted from GBIF. These species had at least 10 HPAI cases from 2021 to 2025.
